# Protein Language Model Predicts Mutation Pathogenicity and Clinical Prognosis

**DOI:** 10.1101/2022.09.30.510294

**Authors:** Xiangling Liu, Xinyu Yang, Linkun Ouyang, Guibing Guo, Jin Su, Ruibin Xi, Ke Yuan, Fajie Yuan

**Affiliations:** Northeastern University; University of Glasgow; Peking University; Westlake University

**Author notes:** Equal Contribution. Work done when Xiangling was a visiting student at Westlake University.

## Abstract

Accurately predicting the effects of mutations in cancer has the potential to improve existing treatments and identify novel therapeutic targets. In this paper, we evidence for the first time that the large-scale pre-trained protein language models (PPLMs) are zero-shot predictors for two *clinically* relevant tasks: identifying disease-causing mutations and predicting patient survival rate. Then we benchmark a series of state-of-the-art (SOTA) PPLMs on 2279 protein variants across 20 cancer-related genes. Our empirical results show that the PPLMs outperform the SOTA baseline, EVE [1], trained on multiple sequence alignment (MSA) data. We also demonstrate that the evolutionary index score, generated from the PPLM’s softmax layer, is good indicator for both mutation pathogenicity and patient survival rate. Our paper has taken a key step toward the clinical utility of large-scale PPLMs.

## 1 Introduction

Whether a mutation causes disease, known as the pathogenicity of a mutation, is a fundamental question in modern genomics. Traditional methods of predicting pathogenicity rely on mutation frequency and clinical evidence in the literature [2]. However, only less than 2% of mutations currently have known functional interpretation [3, 4, 5], leaving most variants have yet to be identified with clinical consequences. Conventional supervised learning models have been tested for this task [6, 7, 8, 9, 10, 11, 12], but the accuracy of these methods remains limited [13].

Recent advances in large-scale language models, such as GPT [14] and BERT, [15] have motivated powerful protein language models by leveraging the similarity between protein sequences and sentences. Among them, the BERT-based pre-trained protein language model (PPLM), a.k.a. ESM-1b [16], trained on 250 million protein sequences in UniRef50 [17], can capture the physical properties of amino acids. The newer version, ESMFold [18], equipped with a much larger PPLM, called ESM2, can predict protein structures with comparable accuracy with AlphaFold [19]. On the other side, a recent study showed [20] that the Evoformer-based PPLM module in AlphaFold2 couldn’t predict protein mutation effects well, perhaps because AlphaFold2 were trained with less fewer protein sequences than ESM1b. A bidirectional LSTM model has been shown to predict whether viruses can escape attacks from hosts’ immune systems [21]. Another similar model is the GPT2-based ProGen2 [22]. Those models can make predictions using a single protein sequence.

There is another direction of research that leverages sequences of homologous proteins. Examples include BERT-based ESM-MSA [23] and VAE-based DeepSequence [24] model. While these models have been tested for predicting mutation effects acrosss a variety of deep mutational scanning experiments, focused assessments in cancer-related mutations are still lacking. More importantly, whether these PPLMs can predict clinically relevant properties (i.e. prognosis) of the cancer patients carrying these mutations remains unknown.

In this paper, we present a systematic benchmark study of state-of-the-art protein language models in predicting pathogenic mutations in cancer diver genes. The models include alignment-based methods, i.e. EVE [1] and ESM-MSA [23], and alignment free approaches, including ESM-1, ESM-1b [16], ESM-1v [25], ESM-2 [18], and ProGen2 [22]. We examine whether these models can learn functional changes caused by mutations in amino acid sequences and identify high-risk mutant cancer proteins. We further test representations learned from these PPLM in a cox-regression framework for progression-free survival prediction. Evaluated on 10,248 patients from The Cancer Genome Atlas (TCGA), the winning model, ESM-1b, can achieve statically significant separation of high and low risk patients in six cancer types, while traditional methods are known to struggle.

## 2 Method and Analysis

### 2.1 Zero-shot Prediction of Pathogenic Mutations using Pre-Trained Protein Language Models

Here, we lay out a zero-shot pathogenic mutation prediction task for pre-trained protein language models. As illustrated in Figure 1, for each each mutation, the model will take the entire protein sequence as input. The output is the probability of sequence being pathogenic or benign. These probabilities are assignment probabilities obtained from fitting a two-component Gaussian mixture to a measurement called evolutionary index. Evolutionary index (EI) is the negative log ratio between the probability of observing the mutant sequence and the probability of observing the wild-type sequence (i.e. sequence without mutation) [24, 1]. The EI has been shown to be an intuitive score to reflect a protein language model’s ability to capture information encoded in amino acid sequences [1, 21]. A higher value in EI indicates stronger deviation of the mutant sequence from the wild-type sequence. Therefore, the component with a higher mean value represents a higher chance of being pathogenic. A straightforward classification can be based on the probability of assigning a mutation to the pathogenic cluster as the pathogenicity score.

**Figure 1:**
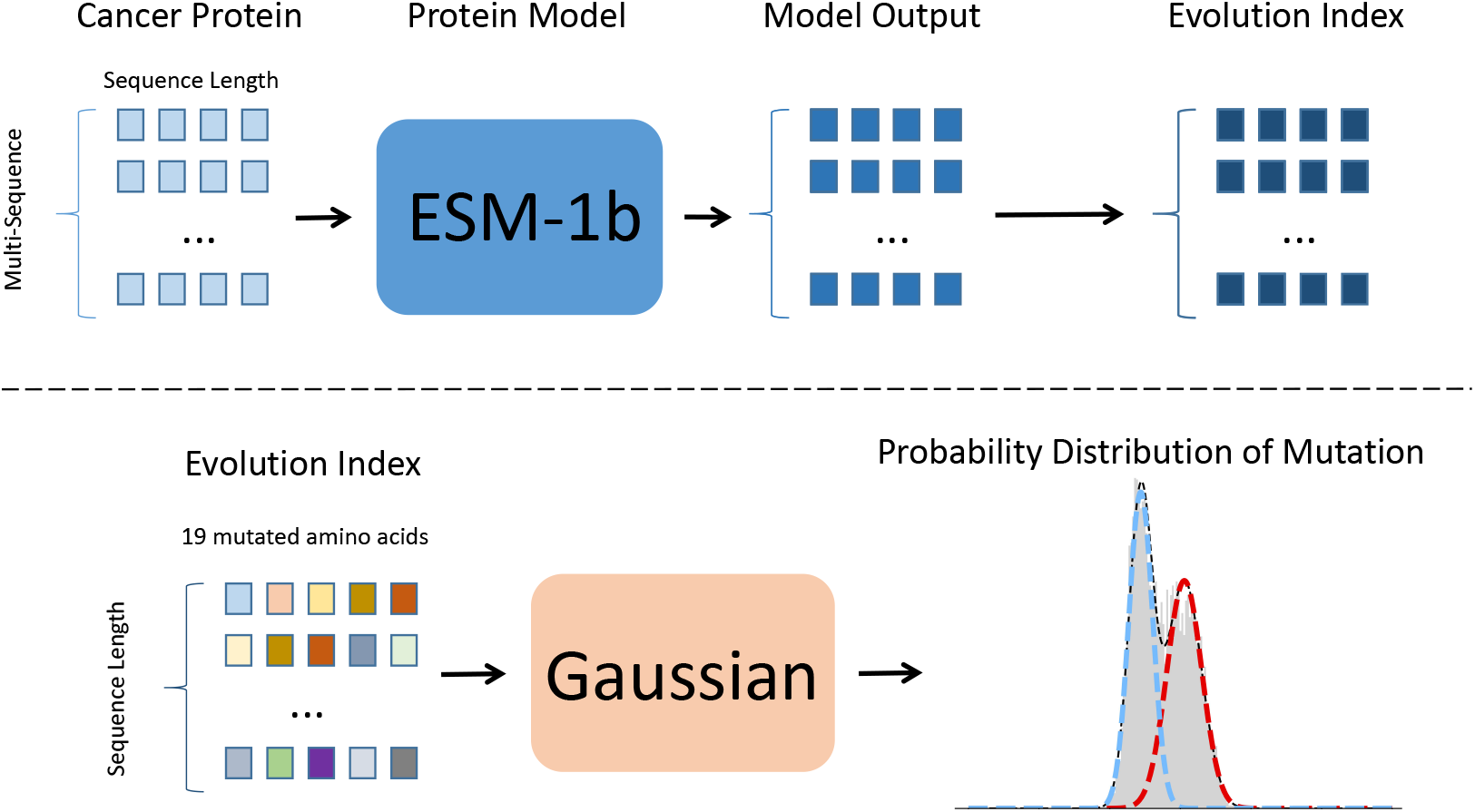
Zero-shot pathogenic mutation prediction framework, using ESM-1b as an example.

The strategy allow us to test pre-trained protein language models solely using their representations, avoiding any confounding factors with further tuning a downstream classifier.

We extracted 20 common cancer proteins from ClinVar [5], which documents and provides supporting evidence for the relationship between variation and phenotype in humans. ClinVar provided the assessment criteria for the clinical significance of variation. ClinVar’s annotations are widely accepted as ground truth. We screened for human cancer proteins, aiming to select as many representative cancer proteins as possible with clinical data. It is important to note that the importation of the protein into the ESM-1b network, the evolutionary index, and the final pathogenicity score were unsupervised and unadulterated with homologous sequence information.

Testing on the ClinVar labels, ESM-1b obtained an AUC of 0.874 and an average ACC of 0.826 (shown in Figure 2), while ESM-1v achieved better results with an AUC of 0.909 (shown in Table 2), outperforming the current leading performance reported [1]. The pathogenic score makes the performance highly explainable. The degree of separation between two Gaussian components directly assesses the strength of the predictive signal captured by ESM-1b. By aligning the score with the genomic position of corresponding mutations, one can further examine the biological implication of the predictions.

**Figure 2:**
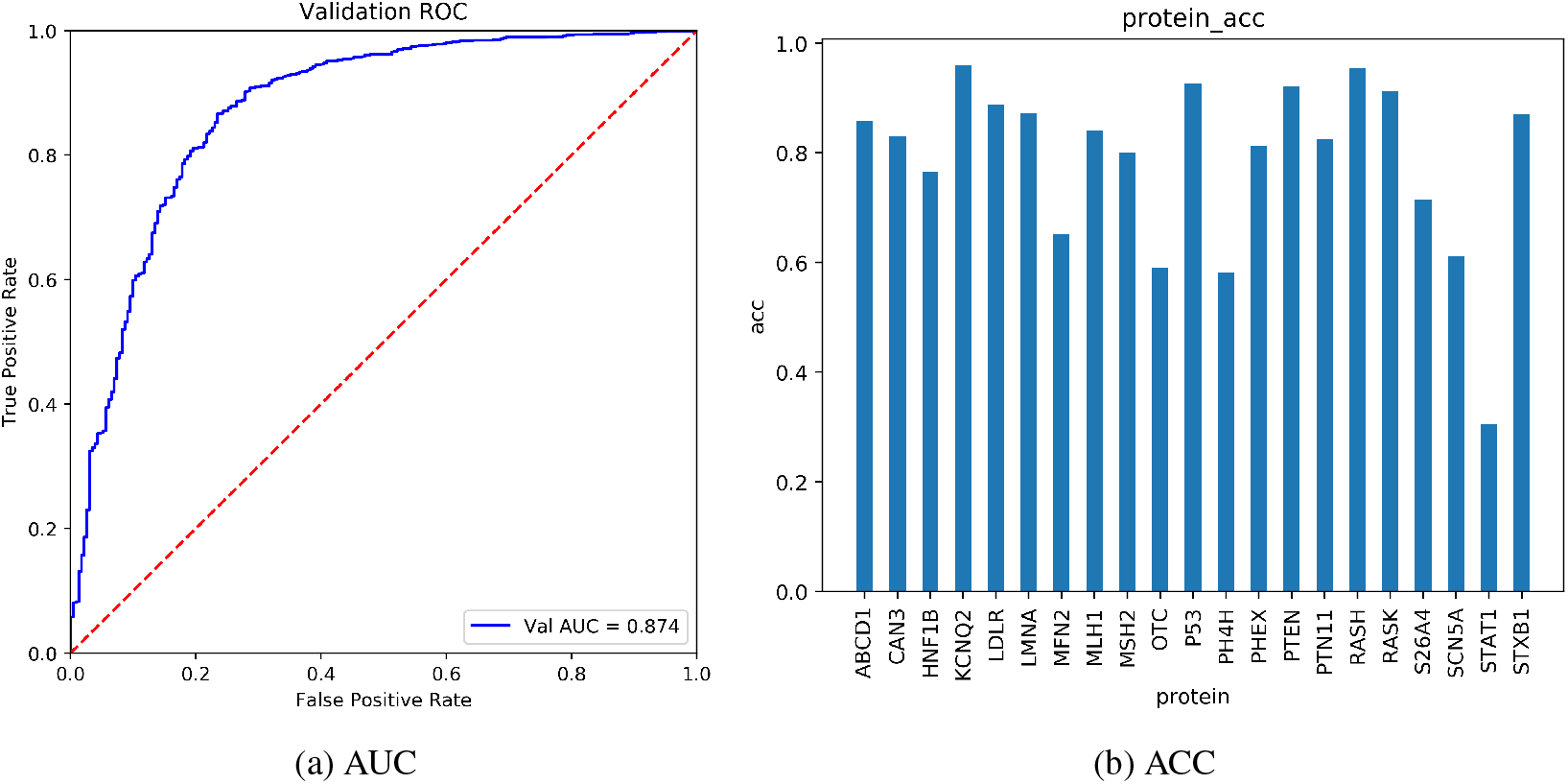
(a) shows the AUC (area under curve) of the pathogenicity scores of 20 common cancer proteins under the Clinvar clinical label, and (b) shows the accuracy of the pathogenicity labels of 20 cancer proteins.

**Figure 3:**
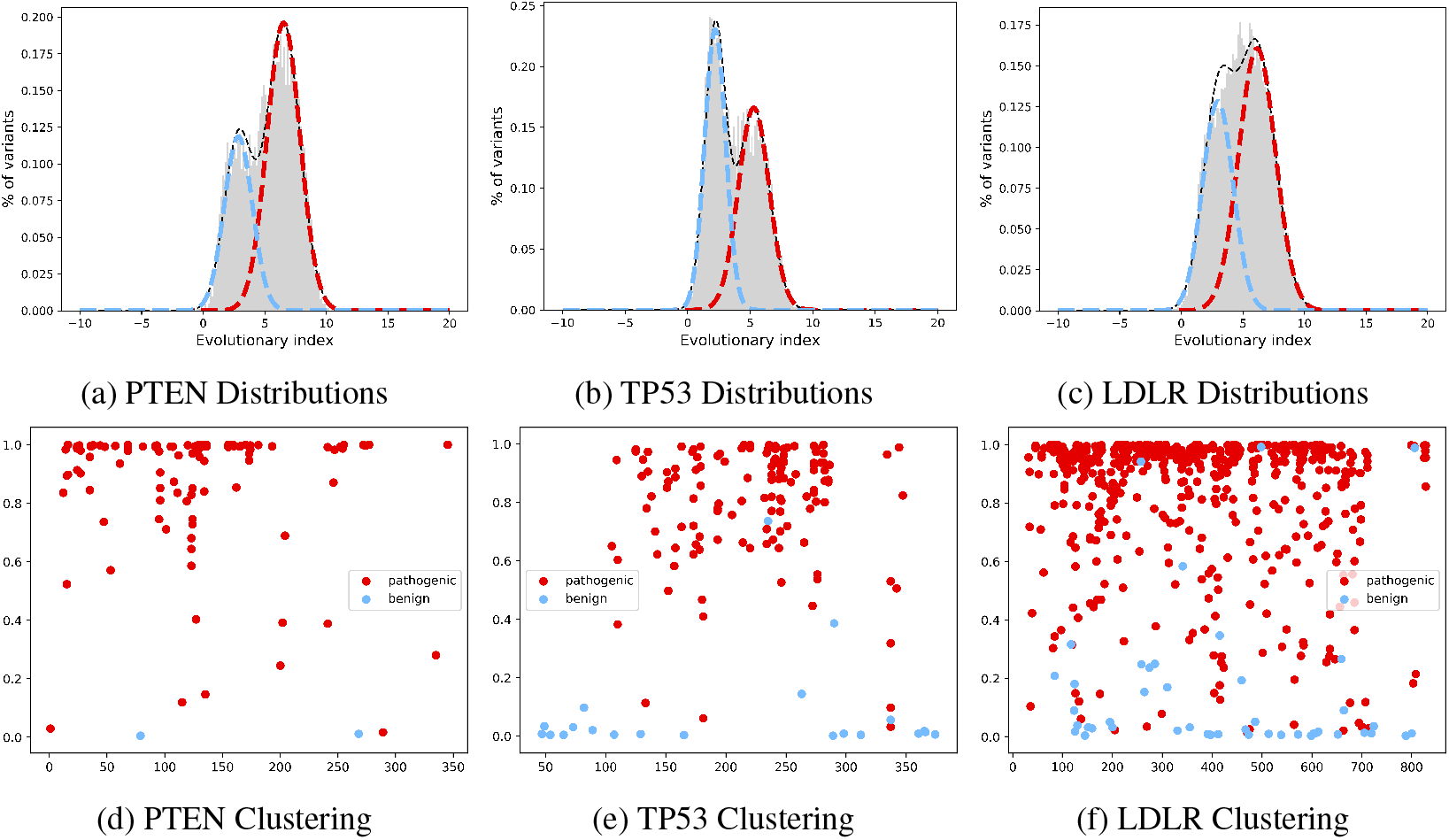
(a), (b), and (c) show the fitting of two-component Gaussian mixture models on evolutionary index scores in PTEN, TP53 and LDLR. There is a clear distinction between benign (blue dashed line) and pathogenic (red dashed line) components. (d), (e), and (f) show the relationship between the pathogenicity score (i.e., the probability of being in the pathogenic component) colour-coded using Clinvar annotations where red denotes pathogenic and blue denotes benign.

**Table 1:** ProGen2 Model.

| Scale | 151M | 764M | 2.7B | 6.4B |
| --- | --- | --- | --- | --- |
| AUC | 0.794 | 0.825 | 0.850 | 0.862 |

**Table 2:** Benchmark protein language models with zero-shot pathogenicity prediction.

| Model | MSA | Params | AUC |  |
| --- | --- | --- | --- | --- |
|  |  |  | Mask Predict | Without Mask |
| EVE | 1 | - | - | 0.887 |
| ESM-1 | 0 | 43M | 0.722 | 0.725 |
|  |  | 85M | 0.780 | 0.769 |
|  |  | 670M | 0.866 | 0.856 |
| ESM-1b | 0 | 650M | 0.706 | 0.874 |
| ESM-1v | 0 | 650M | 0.758 | 0.909 |
| ESM-MSA | 1 | 100M | - | 0.598 |
| ESM-2 | 0 | 8M | 0.695 | 0.701 |
|  |  | 35M | 0.723 | 0.729 |
|  |  | 150M | 0.764 | 0.800 |
|  |  | 650M | 0.827 | 0.822 |
|  |  | 3B | 0.812 | 0.807 |
|  |  | 15B | - | 0.799 |

### 2.2 Benchmark Protein Language Models with Zero-Shot Pathogenicity Prediction

Here, we examine whether pre-trained protein language models can learn functional changes caused by mutations in amino acid sequences and identify high-risk mutant cancer proteins, using the zero-shot prediction task described in the previous subsection. We base our evaluation on six recently proposed large-scale pretrained protein models. The models include alignment-based methods such as EVE [1] and ESM-MSA [23], and alignment free approaches such as ESM-1, ESM-1b [16], ESM-1v [25],ESM-2[18],and ProGen2 [22].

To test whether wild-type context information aid the prediction of mutation pathogenicity, we introduce an additional consideration when testing the model. The input protein sequences follow two settings: one is making the input protein sequence completely visible to the model; the other is we mask amino acid sites of the input protein sequence. When we do not mask, the model learns the probability of this amino acid mutating into other amino acids based on the context information of the mutation site and the wild-type. When masking, the model learns what the masked amino acid should be based on the context information of the mutation site. In our experiment results, we found that masking amino acid sites on the input protein sequences has worse performance on the model’s prediction ability (without masked ESM-1b: AUC=0.874; masked ESM-1b: AUC=0.706, as shown in Table 2).

At the same time, we found that models trained without MSA, such as ESM-1b and ESM-1v, were no worse or even better than models trained with MSA, such as EVE (ESM-1v: AUC=0.909; EVE: AUC=0.887). This result suggests that biological information learned from large-scale protein databases was richer than that from specialized homologous sequences, with the advantage of much less computational time. At the same time, we found that the larger the training dataset, the better the model’s prediction ability.

We also evaluated the performance among models of different scales. We found that the size of a pre-trained protein model is not proportional to its ability to learn the pathogenicity of protein mutations (With the increase in model scale, the AUC of ESM-1 was 0.725, 0.769, 0.874, and 0.856, respectively, shown in Figure 4, and the same as ESM-2). The Zero-shot pathogenicity prediction performance of ESM-1 and ESM-2 does not improve with the increase in model scale; it reaches a peak (at 650M parameters) and then gradually decreases. We also tested the generative model ProGen2, as shown in Table 1. We found that the generative model could also perform a good pathogenicity prediction, and the best 6.4B model reached AUC = 0.862.

**Figure 4:**
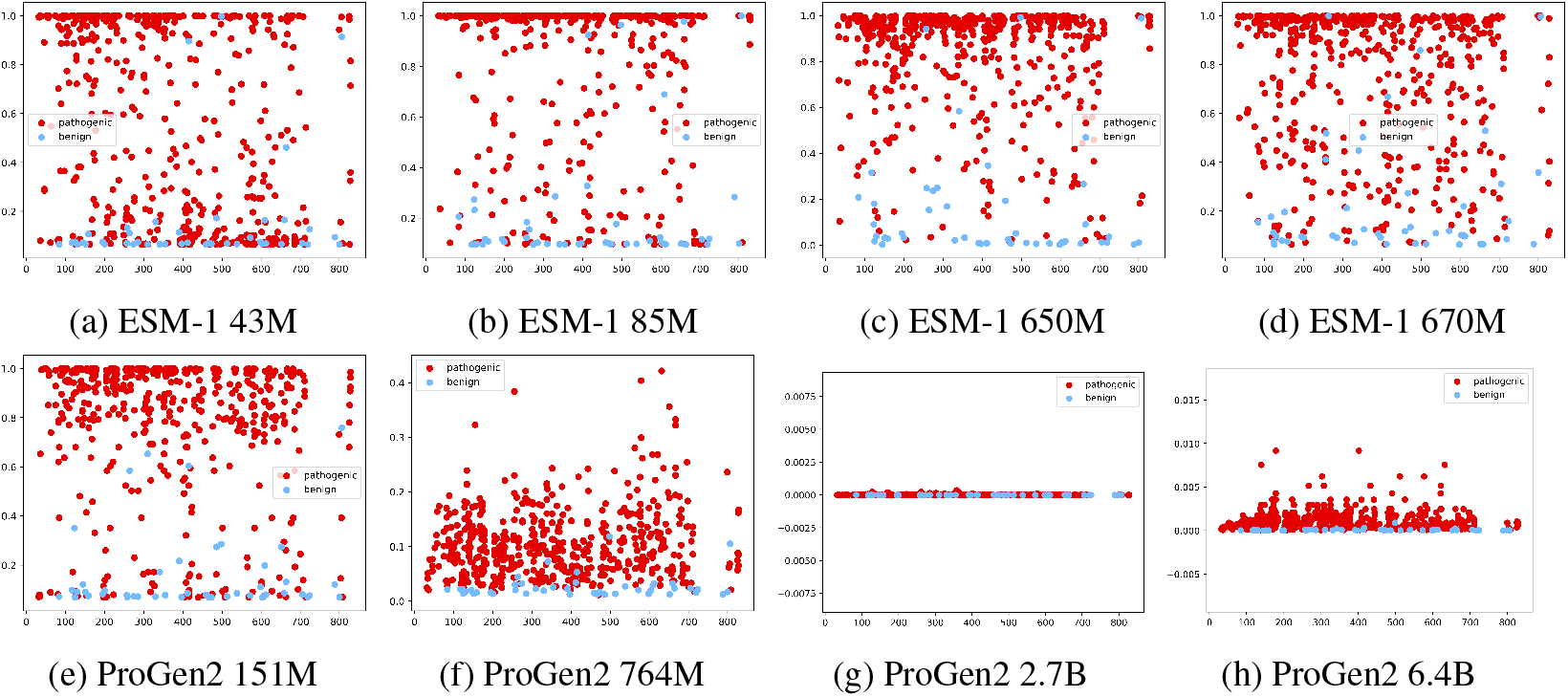
Variation in pathogenicity score of mutations in LDLR with different number of parameters in ESM-1 and ProGen2.

### 2.3 Predict the Clinical Prognosis of Cancer Patients

To investigate the clinical utility of pre-trained protein language models, we examined the prognostic value of the evolutionary index computed using ESM-1b on real-world cancer patients’ data. TCGA is one of the largest datasets from which matched tumour genomic sequencing and clinical outcome data are publicly available [26]. We selected 412 cancer driver genes with suitable protein canonical sequences out of the 579 Tier-1 genes in the COSMIC (Catalogues of somatic mutation in cancer) [27] cancer gene census. EI was estimated by the ESM-1b model for each protein sequence in our driver gene list for each patient.

We performed multivariate Cox proportional-hazards regression (stratified by gender and age) on proteins’ EIs for 10,248 patients across different TCGA cohorts with progression-free interval (PFI) as the adverse outcome. For the 13 cancer types for which our framework was effective, we found that high EI values of specific proteins significantly contribute to better/worse survival (Figure 5a). For example, the high EI value of SMAD4 and FLNA proteins in colorectal cancer (COAD) showed significant evidence of increased patient hazard risk (p<0.01, log-rank test). In contrast, the high EI value of JAK3 protein contributed to a lower hazard risk (p<0.01, log-rank test) in lung cancer (LUAD).

**Figure 5:**
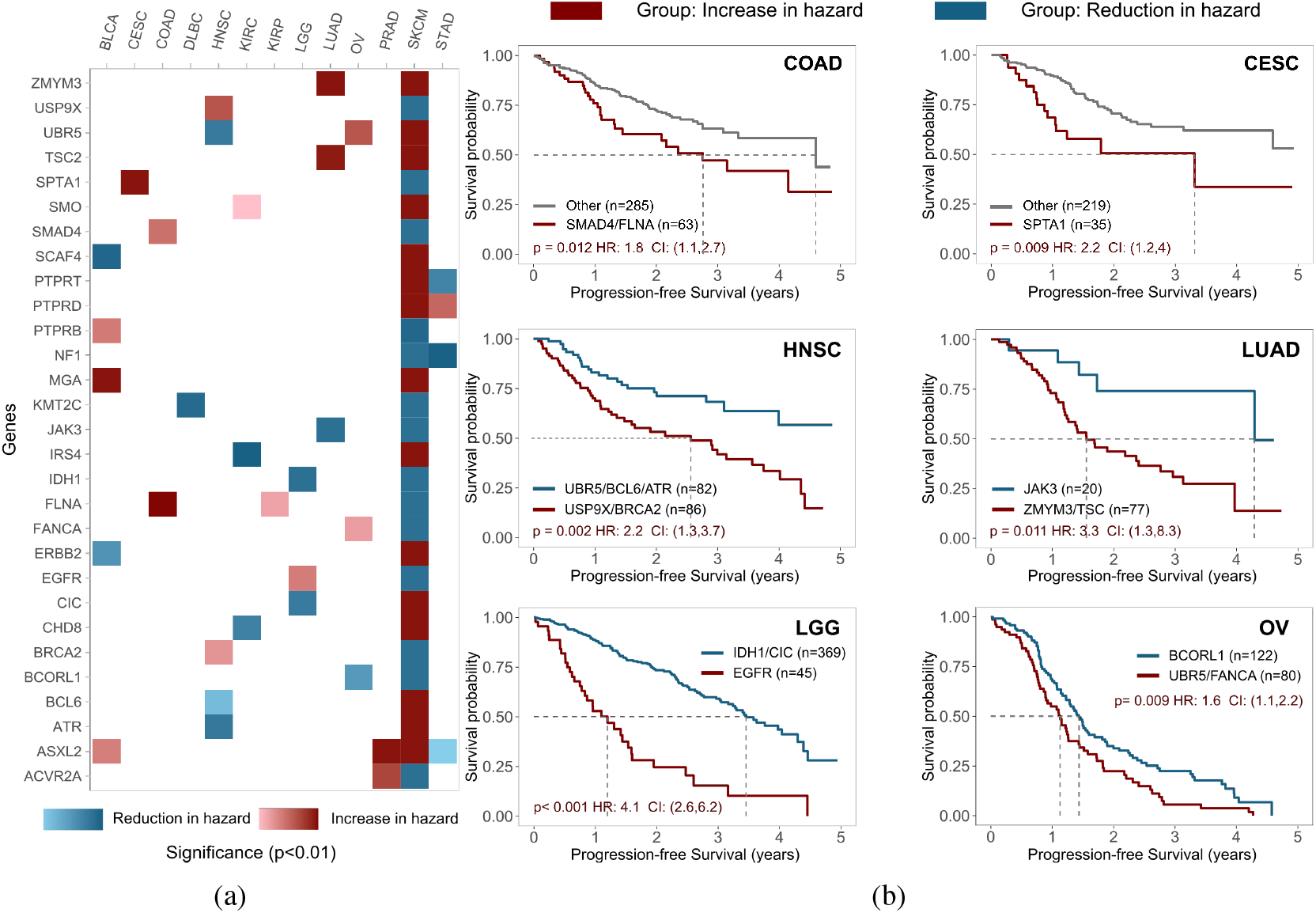
a) Genes with the significant prognostic power across tumours, suggested by multivariate Cox proportional-hazards regression; values in each cell denote the p-value of the log-rank test for corresponding driver genes. b) Kaplan–Meier curves for patient subgroups.

Furthermore, we stratified patients into the hazard-increase and hazard-reduction groups based on whether EI>0 for specific proteins suggested by Cox regression for each cancer type. We drew Kaplan-Meier(KM) curves and tested the survival difference between the two groups with estimated hazard ratios(HR) and p values of the log-rank test. We observed significant difference between hazard-increase and hazard-reduction/other group across 6 cancer types (COAD: p=0.012,HR=1.8,CI=1.1-2.7; CESC: p=0.009,HR=2.2,CI=1.2-4; HNSC: p=0.002,HR=2.2,CI=1.3-3.7; LUAD: p=0.011,HR=3.3,CI=1.3-8.3; LGG: p<0.001,HR=4.1,CI=2.6-6.2; OV: p=0.009,HR=1.6,CI=1.1-2.2; Figure 5b).

These results demonstrate that the evolutionary index can achieve statistically significant survival prediction in multiple cancer types.

## 3 Conclusions

In this paper, we verified that the large-scale pre-trained protein language models can efficiently and accurately predict the effect of mutations in cancer driver genes. Comparable results were obtained with models learned from homologous sequences and those learned from single sequences.

We propose a systematic benchmark based on a zero-short pathogenic mutation prediction task. The experimental results show that BERT-like models such as ESM-1b are better suited to the task than those that rely on generative models. We also found that the size of a pre-trained protein model is not proportional to its performance in predicting pathogenic mutations. This observation aligns with DeepMind’s finding that model performance might drop as the number of model parameters increases, because a large model might be under-trained with limited data [28]. In our case, the complexity and diversity of protein sequences might have been a limiting factor for sufficiently training large models. It is widely hypothesised that existing protein databases only capture a fraction of proteins that exist in living organisms. Finally, we demonstrated the prognostic value of protein language model in TCGA cohorts. The pathogenicity information captured by pre-trained protein model can separate high and low risk patients in six cancer types, while traditional methods have yet been demonstrated success.

## Acknowledgement

This work is supported by the National Natural Science Foundation of China (No.U21A20427, 61972078), the special funding from the Westlake Center of Synthetic Biology and Integrated Bio-engineering (WE-SynBio), and the Research Center for Industries of the Future (No. WU2022C030).

